# A Mixture Copula Bayesian Network Model for Multimodal Genomic Data

**DOI:** 10.1101/110288

**Authors:** Qingyang Zhang, Xuan Shi

## Abstract

Gaussian Bayesian networks have become a widely used framework to estimate directed associations between joint Gaussian variables, where the network structure encodes decomposition of multivariate normal density into local terms. However, the resulting estimates can be inaccurate when normality assumption is moderately or severely violated, making it unsuitable to deal with recent genomic data such as the Cancer Genome Atlas data. In the present paper, we propose a mixture copula Bayesian network model which provides great flexibility in modeling non-Gaussian and multimodal data for causal inference. The parameters in mixture copula functions can be efficiently estimated by a routine Expectation-Maximization algorithm. A heuristic search algorithm based on Bayesian information criterion is developed to estimate the network structure, and prediction can be further improved by the best-scoring network out of multiple predictions from random initial values. Our method outperforms Gaussian Bayesian networks and regular copula Bayesian networks in terms of modeling flexibility and prediction accuracy, as demonstrated using a cell signaling dataset. We apply the proposed methods to the Cancer Genome Atlas data to study the genetic and epigenetic pathways that underlie serous ovarian cancer.

## 1 Introduction

In recent years, there has been considerable interest in estimating causal relationships between random variables in a graphical framework. Among several types of graphical models, Bayesian networks (BN) or equivalently, probability-weighted directed acyclic graphs (DAG) have received the most attention due to their simplicity and flexibility in modeling directed associations in the domain [1, 2, 3, 4]. The associations between d random variables can be summarized by a graph 𝒢 *=* (*V*,*E*) in which *V* = {*X_i_|i =* 1,2,…,*d*} represents the set of variables and *E* ⊂ *V* × *V* represents the dependency between variables. Under the acyclicity and Markov assumptions, the joint likelihood function of (*X*_1_,…,*X_d_*) in a BN has the following simple form based on the conditional densities:

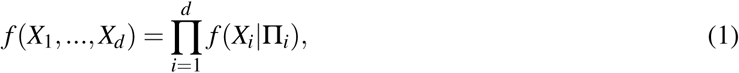

where Π_*i*_ denotes the parent set of *X_i_*, i.e., Π_*i*_ = {*X_j_|X_j_* → *X_i_,X_j_* ∈ *V* \ {*X_i_*}} (Π_*i*_ can be empty).

The two most popular BN models are Gaussian Bayesian network (GBN) model [1] and multinomial Bayesian network (MBN) model [5], for continuous variables and discrete variables respectively. MBN models suffer from super-exponentially increasing number of parameters, therefore can only estimate small-scale networks in practice [5]. To deal with networks with relatively large number of nodes, GBN models have been commonly used due to their simple setup and efficient estimation. However, GBN models may fail to identify the true causalities when the joint distribution of interest is far from multivariate normal, for example, when the underlying distribution is bimodal or multimodal. To tackle the problem of non-normality, several new BN models have been developed, for instance, the logistic Bayesian network by Zhang et al. [4] which discretizes all the continuous variables to fit a multi-category logit model. Considerable work has also been done in nonparametric and semiparametric estimation of the BN structure. For instance, Voorman el al. [6] proposed the following nonparametric model to deal with non-normality issue:

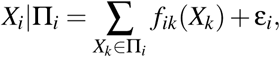

where the *f_ik_* (·) lies in some function space **ℱ**. The model by Voorman et al. focuses on estimating the conditional mean *E*(*X_i_*|Π*i*). It is essentially a generalized additive model without assuming the independence between ε_i_ and *f_ik_*(·). However, this method relies on a known causal ordering of the true network which is unavailable in most cases.

In 2010, Elidan [7] introduced an innovative copula Bayesian network (CBN), a marriage between copula functions and graphical models, which extends conventional BN models to a more flexible framework. A CBN model constructs multivariate distribution with univariate marginals and a copula function C that links these marginals. In general, one can estimate marginals using parametric or non-parametric approach, and then use a small number of parameters to capture the dependence structure. However, as we shall see in a real data set (Section 4), the regular copula functions such as Gaussian copula may not be able to accurately depict multimodal joint distributions. In addition, the CBN model is subject to the choice of copula function for each local term. Motivated by Elidan’s work, we extend the regular copula Bayesian networks to a mixture copula Bayesian network (MCBN) using finite mixture models, to better deal with non-normality, multimodality and heavy tails that are commonly seen in current massive genomic data. The parameters in a MCBN model can be efficiently estimated by a routine EM algorithm. As demonstrated by the real data, the performance of a two-component Gaussian MCBN is generally promising, and our model achieves reasonable accuracy in identifying the true edges in a sparse causal network.

The rest of this paper is organized as follows: In Section 2, we review Elidan’s CBN model, and introduce the proposed MCBN model using a two-component Gaussian mixture for illustration. In Section 3, we present a heuristic local search approach combined with a routine EM algorithm for graph structure estimation, as well as the best-scoring network out of multiple predictions with random initial values. The comparison of three BN models is carried out over a cell signaling data set in Section 4. The new model is applied to the Cancer Genome Atlas (TCGA) data for serous ovarian cancer in Section 5. We discuss and conclude this paper in Sections 6 and 7.

## 2 Method

### 2.1 Copula and Elidan’s Copula Bayesian network

Unless otherwise stated, we use 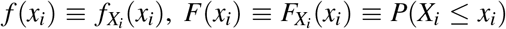 as the marginals, and similarly for multivariate density *f* (**x**) = *f*^**x**^(**x**). The formal definition of copula function is given below:

#### Definition 1.

Let (*X*_1_, *X*_2_, …, *X_d_*) be a vector of continuous random variables and (*F*(*x*_1_), *F*(*x*_2_), …, *F*(*x_d_*)) be the marginal distribution functions. The copula function of (*X*_1_, *X*_2_, …, *X_d_*), *C*: [0, 1]^*d*^ → [0, 1], is defined as the cumulative distribution function of (*F*(*X*_1_), *F*(*X*_2_), …, *F(X_d_*)):

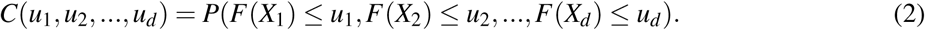

By definition, a copula function is a multivariate distribution function where the marginals are uniform. By choosing an appropriate copula, one can generate multivariate distribution of any complex form. In practice, one can completely separate the choice of marginals and the choice of dependency patterns between random variables. Sklar’s Theorem below guarantees that any multivariate distribution can be expressed with univariate marginals and a copula function which links these variables:

#### Theorem 1.

Let *F* (*x*_1_, *x*_2_,…, *x_d_*) be a multivariate distribution over real-valued d-dimension random vectors, then there exists a copula function that satisfies:

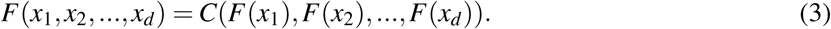

Furthermore, the copula function C is unique when the marginal distribution *F*(*x_i_*) is continuous for *i* ∈ {1, 2, …, *d*}.

By taking the first derivative for both sides of Equation (3), we can derive the copula density function defined as 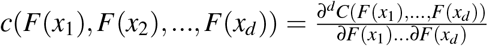. The copula density is simply a ratio between the joint density and the product of all the marginals:

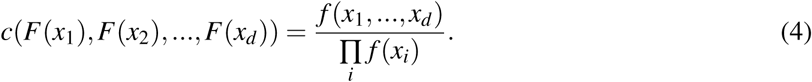

An immediate consequence of Equation (4) is that *c*(*F*(*x*_1_), *F* (*x*_2_), …, *F* (*x_d_*)) = 1 if and only if *X*_1_, …, *X_d_* are independent. For a subset of variables (*Y, X*_1_, …, *X_p_*), as 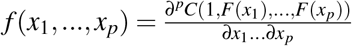, the conditional density *f* (*y*|*x*_1_, …, *x_p_*) can be expressed as follows:

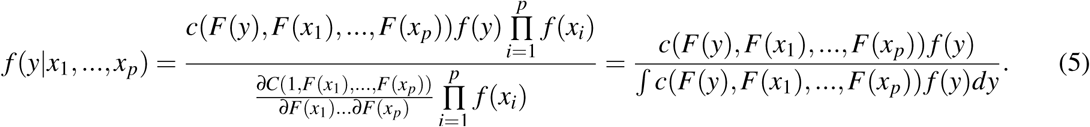

Motivated by Equations (1) and (5), Elidan proposed a copula Bayesian network based on the following local density:

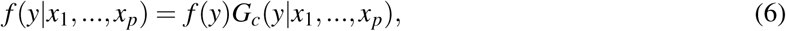

where 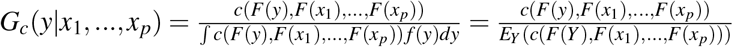.

By Equation (6), we have the following decomposition for the joint density of variables in a Bayesian network:

#### Theorem 2.

Let (*X*_1_, …, *X_d_*) be *d* random variables (nodes) in a Bayesian network, and π_*i*_ = {*x_j_*|*X_j_* ∈ Π_*i*_}. The joint density can be represented as follows:

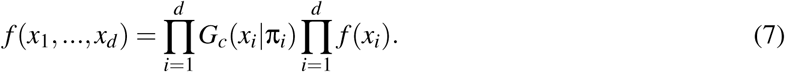

Although the construction of local copulas can significantly reduce the complexity of the structure learning, choosing an appropriate copula for each local term *G_c_*(*x_i_*|π_*i*_) is essential. Elidan suggested a small set of pre-selected copula functions (or copula families) such as Gaussian copula, Frank’s copula, Ali-Mikhail-Haq (AMH) copula and Gumbel-Barnett (GB) copula. However, as we will discuss in Section 4, these regular copula functions might be inadequate to model the complex dependence structure. To this end, we extend the copula Bayesian network to a more flexible framework using finite mixture model.

### 2.2 A Mixture Copula Bayesian Network

For illustration purpose, we limit ourselves to Gaussian MCBN, but other mixture models such as Gamma mixture and Beta mixture models can be adapted similarly. The K-component Gaussian mixture copula for variables (*Y*, *X*_1_, …, *X_p_*) can be formulated as follows:

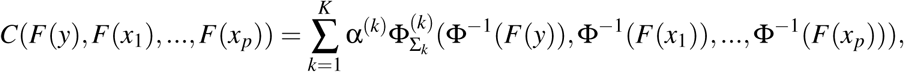

where *α*^(*k*)^ and 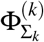 denote the weight and cumulative distribution function (CDF) of the kth Gaussian component respectively, and Φ^−1^(·) represents the quantile function of *N*(0, 1). The corresponding copula density can be obtained immediately:

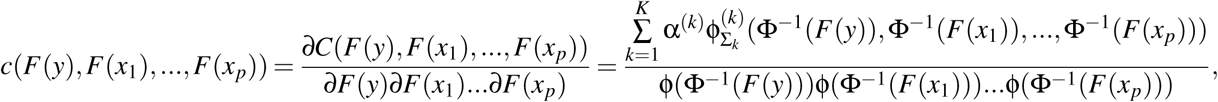

where ϕ(·) represents the standard normal density function.

The Gaussian MCBN model above takes advantage of finite mixture model to better fit the bimodal and multimodal distributions. Similar as in the Elidan’s copula Bayesian network, the marginals should be estimated prior to fitting the mixture copula, with either parametric or nonparametric method. We can, for example, fit the marginals using parametric or nonparametric method, then transform (*y*, *x*_1_, …, *x_p_*) to (*F*(*y*),*F*(*x*_1_), …, *F*(*x_p_*)) using the fitted CDF functions. The transformed values will be used for estimating the copula function. Based on the estimated mixture copula for each local term in BN, we can calculate the joint likelihood by Equation (7).

## 3 Graph estimation using EM and local search algorithms

### 3.1 EM algorithm for finite Gaussian mixture

In this part, we introduce the EM algorithm to estimate the mixture copula for each local term *G_c_*(*x_i_*|π_*i*_).

For a given variable *X_i_* and its parent set Π_*i*_, the regular k-means algorithm can provide warm starts for the mean vector ***μ***_*k*_ (of dimension | Π_*i*_ | +1) and the covariance matrix **Σ**_*k*_ (of dimension (| Π_*i*_ | +1) × (| Π_*i*_ | +1)) for each mixture component, as well as the mixing rate *α*^(*k*)^. Let 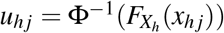 and ***u***_*j*_ = {*u_hj_*}, where *x_hj_* is the observed value for variable *X_h_* and sample *j*, *X_h_ ∈* {*X_i_*, Π_*i*_ }, *j* = 1, 2, …, *N*. Let ***Z*** =(*z*_1_,…,*Z_N_*) be the vector of indicators for the membership of each sample (mutually exclusive and exhaustive), i.e., *α*^(*k*)^ *= p(z_j_ = k), j =* 1, …, N and 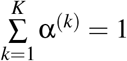. Denote Θ_*k*_ = (***μ***_*k*_, **Σ**_k_) and Θ = {Θ_*k*_}, the EM algorithm with missing information ***Z*** can be implemented as follows:

- **E Step:** Given current estimate of all the parameters (α^(*k*)^, **Θ**), we compute the weighted membership as follows:

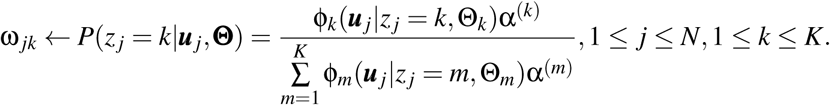
- **M Step:** Use data ***u***_*j*_ and membership weights to update all the parameters:

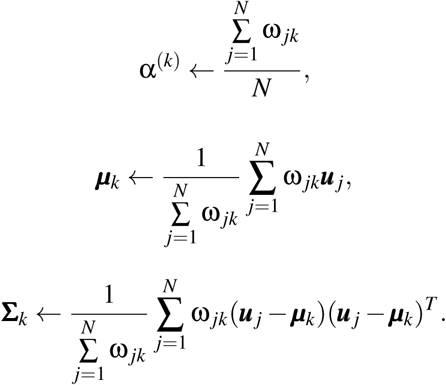

Given an estimate of the graph structure *𝒢* and the parameters scriptthetta Θ^(𝒢^), the log-likelihood can be written as:

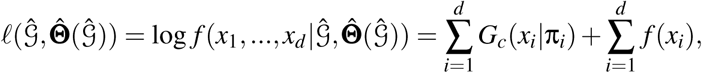

where the denominator of *G_c_*(*x_i_*|π_*i*_), i.e., *E_Xi_*(*c*(*F*(*X_i_*)*,F*(*π_il_*)*, …, F*(*π_ipi_*))) must be evaluated. Here we use notation *p_i_* as the number of parents of *X_i_,* i.e., *p_i_ =* |Π_*i*_|. A simple idea for estimating *G_c_*(*x_i_*|π_*i*_) is to generate a list of Monte Carlo samples 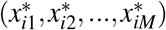 from *f (x_i_),* and by law of large numbers:

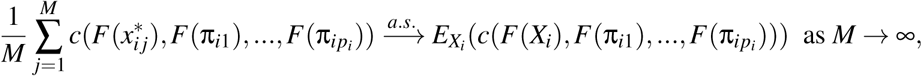

where 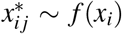. However, it is noteworthy that drawing samples from *f (x_i_)* might be complicated and time-consuming when marginals were estimated with nonparametric method. Further, the likelihood ℓ(𝒢^, Θ^ (𝒢^)) may fail to converge due to the randomness of *G_c_*(*x_i_*|*π_i_*) estimation. Therefore for practical consideration, one can directly use all the observations as samples so that the convergence is guaranteed.

### 3.2 Score-based local search for learning MCBN

In this part, we introduce an efficient heuristic search algorithm based on Bayesian information criterion (BIC) to learn the structure of underlying network 𝒢. The BIC score can be evaluated by the following formula:

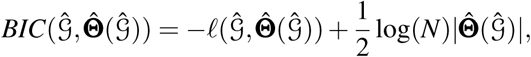

where 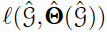 represents log-likelihood function, 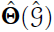 is the set of all the parameters including the mixing rates, mean vectors and covariance matrices of Gaussian components, and 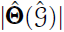 denotes the total number of free parameters in 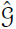. We start from a randomly generated network or empty network, and greedily advances through basic edge operation including addition, deletion and reversal, until BIC score reaches the minimum [7]. Unfortunately, this local search algorithm may easily get trapped in local maximum due to the high dimensionality and non-convexity of the likelihood function, making it impractical to find the global maximum. Enlightened by one of the reviewers, we conducted the heuristic search algorithm for multiple times, each with a random initial value, and the best-scoring network (with minimum BIC score) was returned as the best predicted network.

## 4 Comparison with existing models

In this section, we compare the proposed MCBN model with two existing BN models, including the GBN model and Elidan’s CBN model. We tested the three models using a flow cytometry dataset generated by Sachs et al. [8]. Sachs’ data contains simultaneous measurement on 11 protein and phospholipid components, which was used for elucidating the signaling pathway structure in the cells of human immune system. The known network shown in Figure 1a is a Bayesian Network containing 11 nodes and 20 causal relations. Each causal edge in the network was well validated by experimental intervention, therefore this network structure is often used as the benchmark to assess the accuracy of different directed or undirected graphical models.

**Figure 1:**
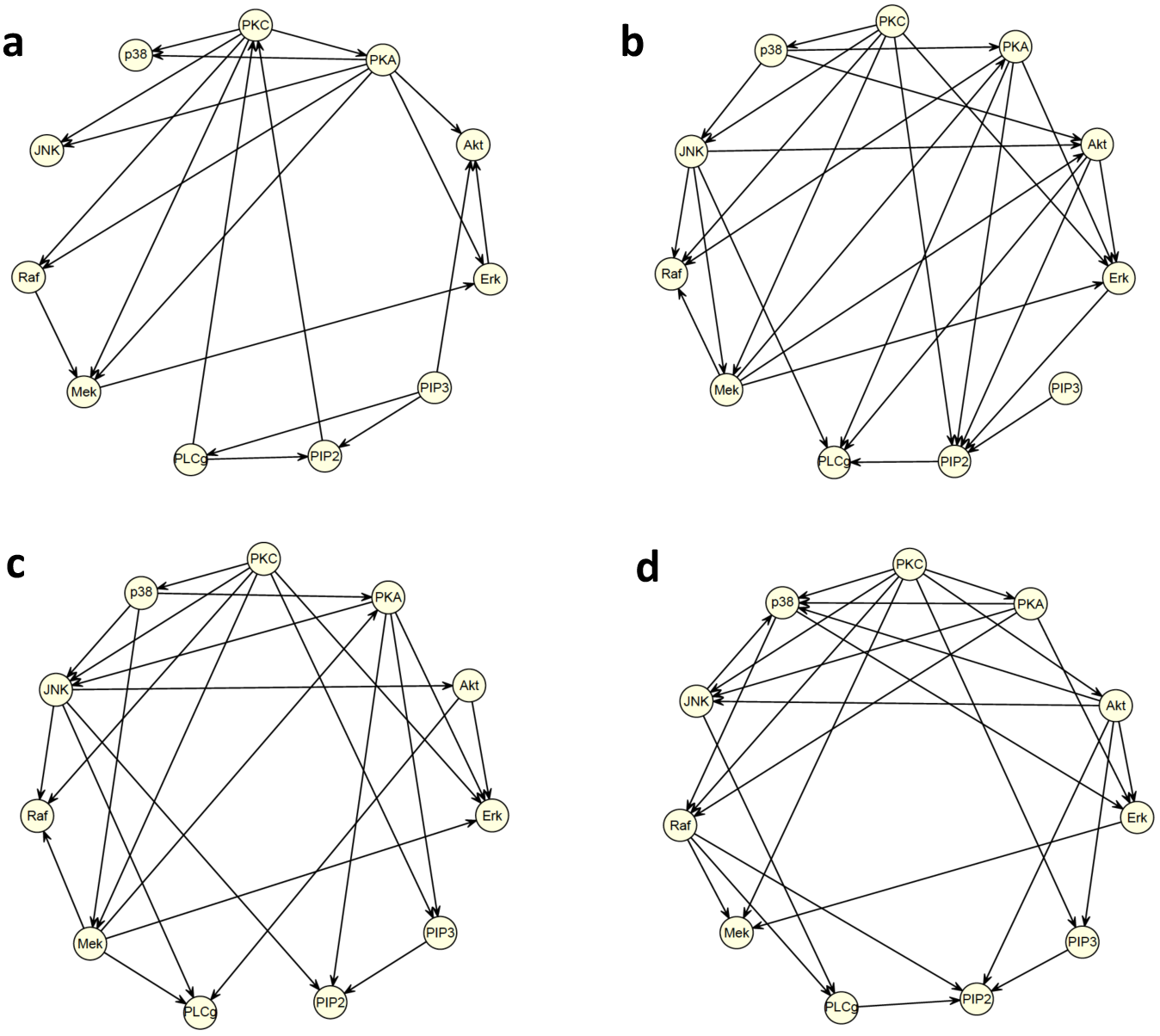
Comparison of three Bayesian network models on Sach’s data: (a) The benchmark network; (b) Network predicted by GBN model; (c) Network predicted by Gaussian CBN model; (d) Network predicted by two-component Gaussian MCBN model.

Sachs’ data has both continuous and discrete versions. In our analysis, we used the continuous data which was log-transformed and normalized by subtracting the mean and dividing by standard deviation. Three BN models were then applied to the preprocessed data for network structure learning, with detailed implementation as follows:

- **GBN:** We considered the linear regression setting, 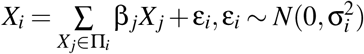, where the graph structure and parameters were estimated by a Blockwise Coordinate Descent (BCD) algorithm proposed by Fu and Zhou [1]. It has been shown that the BCD algorithm outperforms the popular PC algorithm [9] under regular settings. The intervention information was also incorporated in the modeling and a geometric sequence of 100 candidate tuning parameters (λ_1_, …, λ_50_) were predefined (λ_1_ = 0.001, λ_100_ = 1). All the calculations were done using the source code provided by the authors (personal communication).
- **MCBN**: For simplicity of calculation, we considered a two-component Gaussian MCBN. The two-component Gaussian mixture model were also applied to the univariate marginals. Figure 2 shows two examples of fitted marginals for proteins *Art* and *Erk*. We set the maximum number of parental nodes at 5, i.e., max_*i*_ |Π_*i*_| ≤ 5. The local search algorithm with BIC criterion was applied to BN structure learning, starting from an empty network. In the EM estimation of the copula function, we used k-means (*K* = 2) to obtain initial values for all the parameters, and used threshold 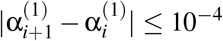 for convergence, where α_*i*+1_^(1)^ and α_*i*_^(1)^ represent the resulting mixing rates in two consecutive EM runs.
- **CBN**: Elidan’s CBN model can be treated as a special case of MCBN model when the copula density function has only one component (Gaussian copula). For the sake of comparison, all the marginals were also fitted using two-component Gaussian mixture. Same threshold as in MCBN was used as convergence criterion of the EM algorithm.

**Figure 2:**
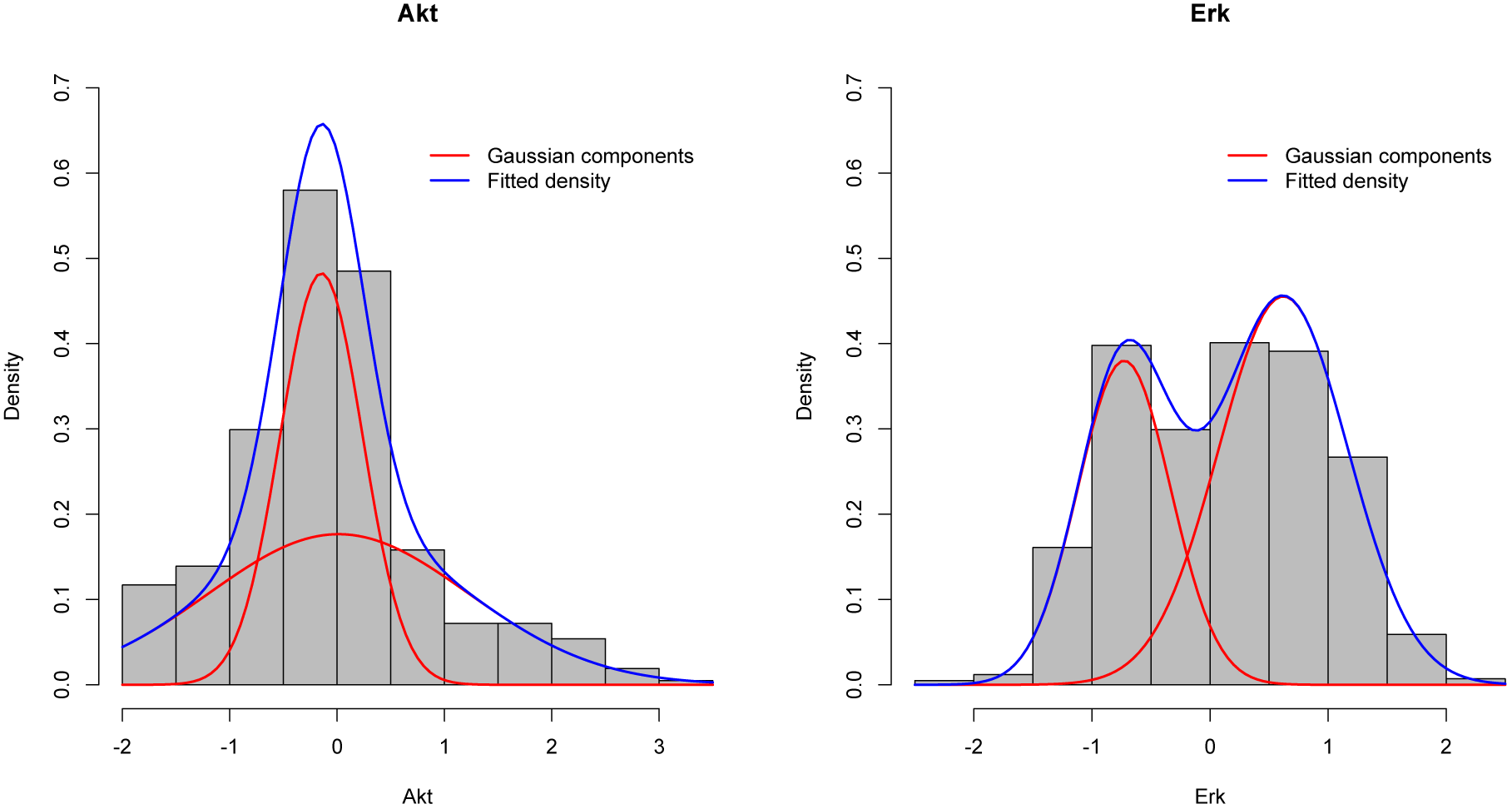
Fitted marginals by a two-component Gaussian mixture for the abundance of proteins *Akt* (left) and *Erk* (right).

The estimated graphs by three different models are shown in Figure 1b-d. Table 1 summarizes true positive rate (TPR), false discovery rate (FDR) as well as running times by the three models (all timing were carried out on a Intel Xeon 3.2GH processor). In this comparison, a predicted edge is considered correct if both connection and direction are correct. It can be seen that the proposed MCBN model achieves significantly higher accuracy than the two existing models in terms of TPR and FDR, but it is more computationally expensive than the two simpler BNs. To further improve prediction, we conducted 100 predictions using random initial networks and obtained the best-scoring network, which contained 25 predicted edges. Out of 20 true edges, 13 were correctly identified in the best-scoring network. Furthermore, we compared different models in capturing the dependency pattern between variables. Figure 3 shows the scatterplot of *Art* and *Erk*, and the plots of simulated samples from three generative models. Compare to other models, the two-component Gaussian MCBN better depicted the multimodal dependency between *Akt* and *Erk*.

**Table 1:**
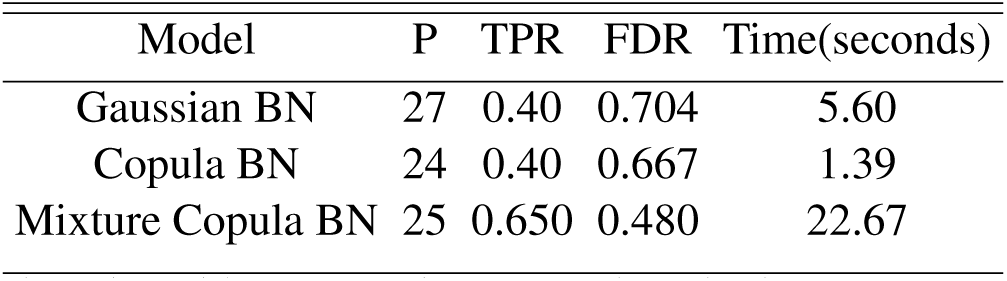
Comparison of three different BN models. Presented in the table are number of predicted edges (P), true positive rate (TPR), false discovery rate (FDR), as well as the CPU time (in seconds) by three different BN models.

| Model | P | TPR | FDR | Time(seconds) |
| --- | --- | --- | --- | --- |
| Gaussian BN | 27 | 0.40 | 0.704 | 5.60 |
| Copula BN | 24 | 0.40 | 0.667 | 1.39 |
| Mixture Copula BN | 25 | 0.650 | 0.480 | 22.67 |

**Figure 3:**
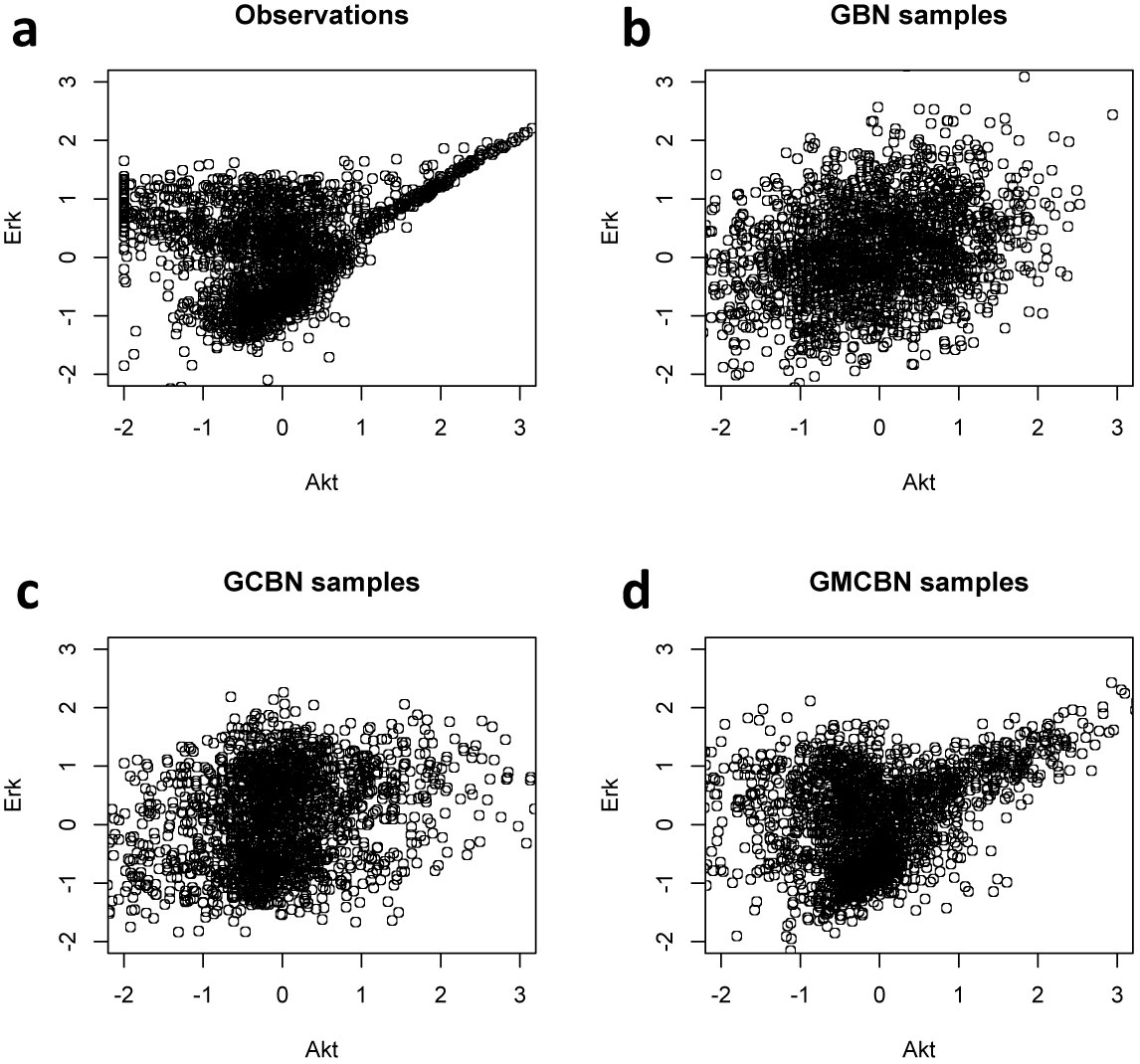
Dependence between proteins *Art* and *Erk:* (a) Observations; (b) Simulated samples from GBN; (c) Simulated samples from Gaussian CBN; (d) Simulated samples from two-component Gaussian MCBN.

To select the most confident edges, we calculated the log-likelihood decrease by removing one edge from the network. We found that an edge giving more likelihood increase has higher probability to be a true edge in the network. For instance, we selected the 10 most confident edges based on the likelihood change, and seven of them turned out to be true edges including *Akt*→*Erk, PKC*→*P38, PIP3*→*PIP2, PKA*→*Raf, PKC*→*JNK, PKC*→*Raf* and *PLCg*→*PIP2.* In addition, we evaluated the performance of our model in predicting the network skeleton (undirected edges). The proposed MCBN was compared with two simple alternatives including Pearson’s correlation and Spearman’s correlation. In this comparison, a predicted edge is considered correct as long as the connection is correct. Figure 4 shows the undirected networks by three approaches, and the TPR/FDR are summarized in Table 2.

**Figure 4:**
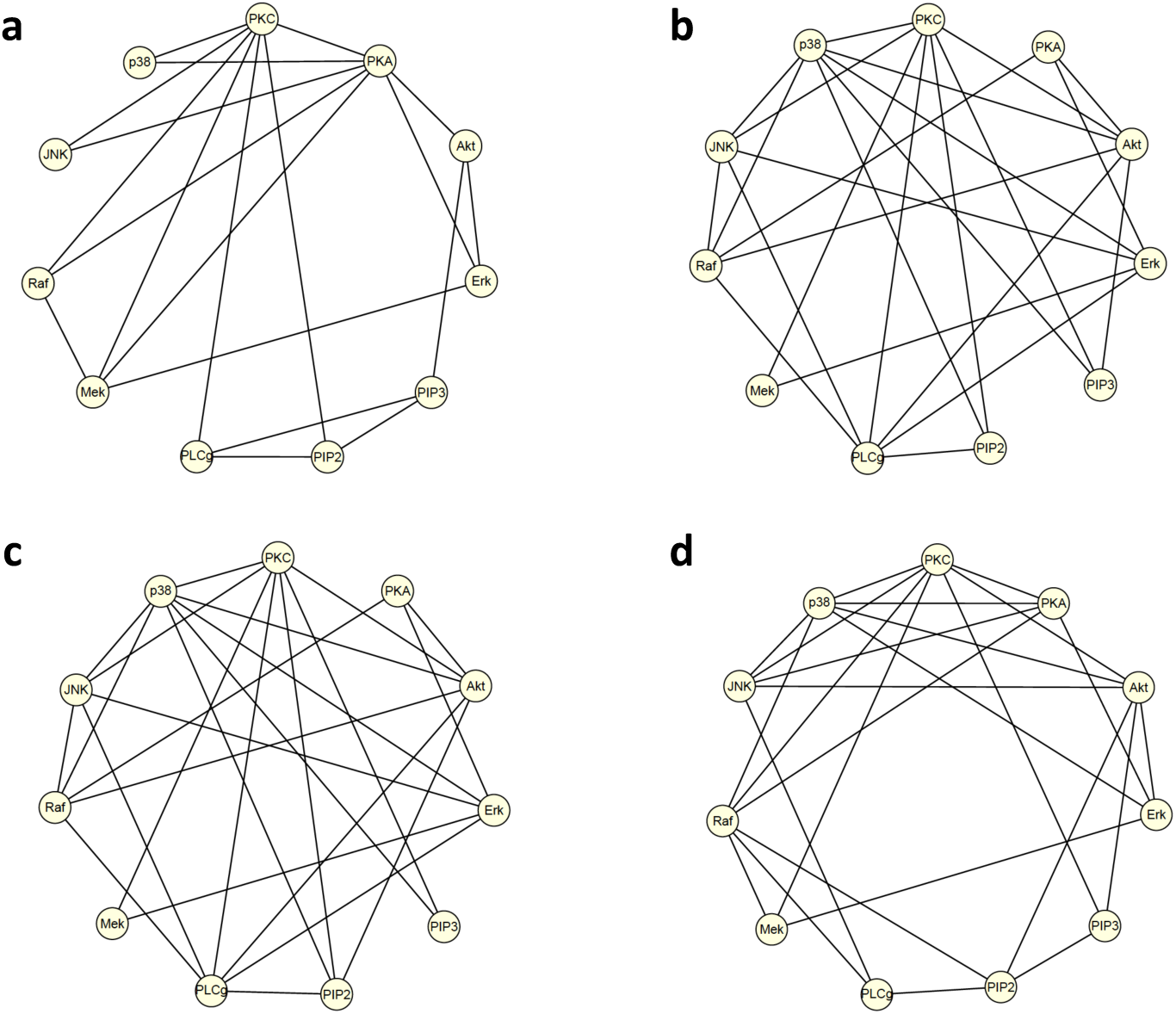
Comparison of three undirected networks: (a) Skeleton of the known network presented in Figure 1a; (b) Network consisted of top 25 edges based on Pearson’s correlation coefficient; (c) Network consisted of top 25 edges based on Spearman’s correlation coefficient; (d) Skeleton of network predicted by MCBN model presented in Figure 1d.

**Table 2:** Comparison with Pearson’s and Spearman’s methods. Presented in the table are number of undirected edges (P), true positive rate (TPR), false discovery rate (FDR) by three different approaches. For Pearson’s and Spearman’s methods, we selected top 25 edges with strongest correlation coefficients.

| Model | P | TPR | FDR |
| --- | --- | --- | --- |
| Pearson's correlation | 25 | 0.55 | 0.56 |
| Spearman's correlation | 25 | 0.50 | 0.60 |
| Mixture Copula BN | 25 | 0.75 | 0.40 |

## 5 Application to TCGA ovarian cancer data

In this section, we applied the proposed MCBN to the Cancer Genome Atlas (TCGA) data [10], to study the interactions between oncomarkers that are associated with serous ovarian cancer. The TCGA data is one of the most comprehensive cancer genomic data sets, with more than 30 cancer types and subtypes which include but not limited to ovarian cancer, breast cancer, lung cancer, brain cancer and liver cancer. The sample sizes range from 50 to 1200 for different cancer types, and each sample is represented by both the molecular profile and clinical information. The molecular profile contains measurements for various types of (epi)genetic factors including gene expression quantification (both microarray and RNAseq), DNA methylation, single nucleotide polymorphism (SNP), copy number variation (CNV), somatic mutation, and microRNA etc. The clinical data provide information such as race, gender, tumor stage, outcome of surgery and resistance to chemotherapy.

The TCGA ovarian cancer data collected 567 tumor samples and 8 organ-specific normal controls. We incorporated three data types into our model including gene expression level, DNA methylation level (on gene promoter region) and CNV. The data were normalized using a quantile normalization method by Balstad et al. [11, 12] to correct the bias due to non-biological causes. In addition, we applied an effective method by Hsu et al. [13] to remove age and batch effects (three age groups are defined as < 40 y.o., [40, 70] y.o., and > 70 y.o.). Hsu’s method is essentially a median-matching and variance-matching strategy. For example, the batch-effect-adjusted gene expression value can be obtained as follows:

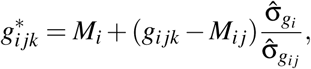

where *g_ijk_* represents the expression level of gene i from batch j and sample k, *M_ij_* denotes the median of *g_ij_ = (g_ij_*_1_*,…, g*ijn), *M_i_* denotes the median of *g_i_ = (g_i_*_1_*,…, g_ij_*), 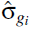 and 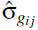 are the standard deviation of *g_i_* and *g_ij_*, respectively.

The set of biomarkers was identified by a stepwise correlation-based feature selector (SCBS) by Zhang et al. [4], which mimics the hierarchy of underlying causal network. The SCBS algorithm starts from selecting the nodes that are strongly associated with the phenotype node and progressively select the nodes that are associated with the selected nodes in previous step. This algorithm is more effective in identifying phenotype-associated nodes, especially those nodes that are indirectly associated with the phenotype. By 3 runs of SCBS, we identified 73 oncomarkers including the expression level of 50 genes, CNV at 15 sites and methylation level at 8 sites. Among the 73 oncomarkers, many were previously reported in the literature including *BRCA1* [10], *BRCA2* [10], *RBI* [14], *PTEN* [15], and *OPCML* [16].

We then fit a MCBN model to study the regulatory relationships between these oncomarkers. The marginals were fitted by a two-component Gaussian mixture (other mixture models can also be used, e.g., Beta-mixture for DNA methylation). Figure 5 and 6 show several examples of the fitted marginals for *TP53* (expression level), *SPARC* (expression level), *BRCA1* (methylation level) and *NOTCH3* (methylation level).

**Figure 5:**
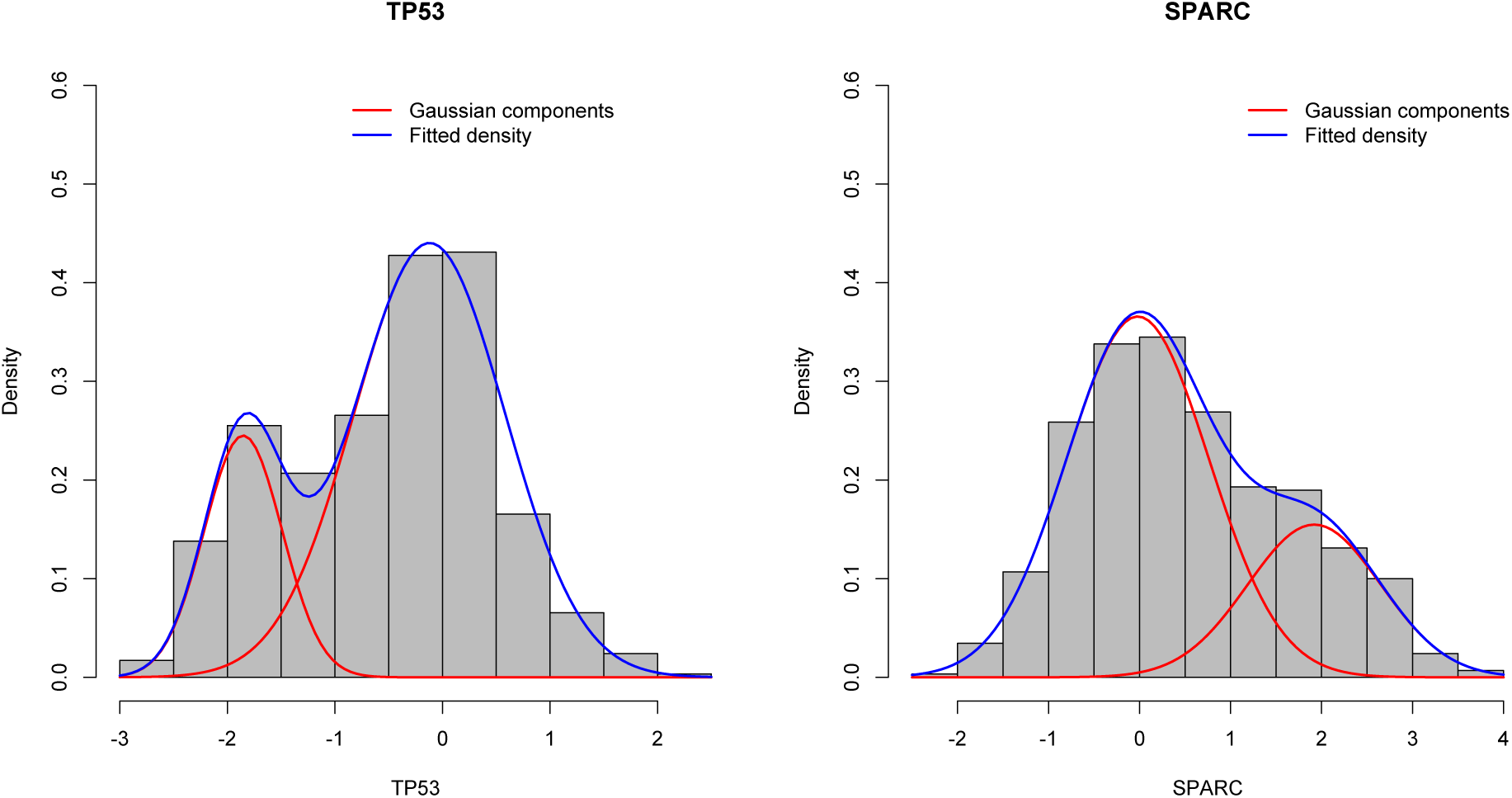
Fitted marginals by a two-component Gaussian mixture for the expression level of gene *TP53* (left) and *SPARC* (right).

**Figure 6:**
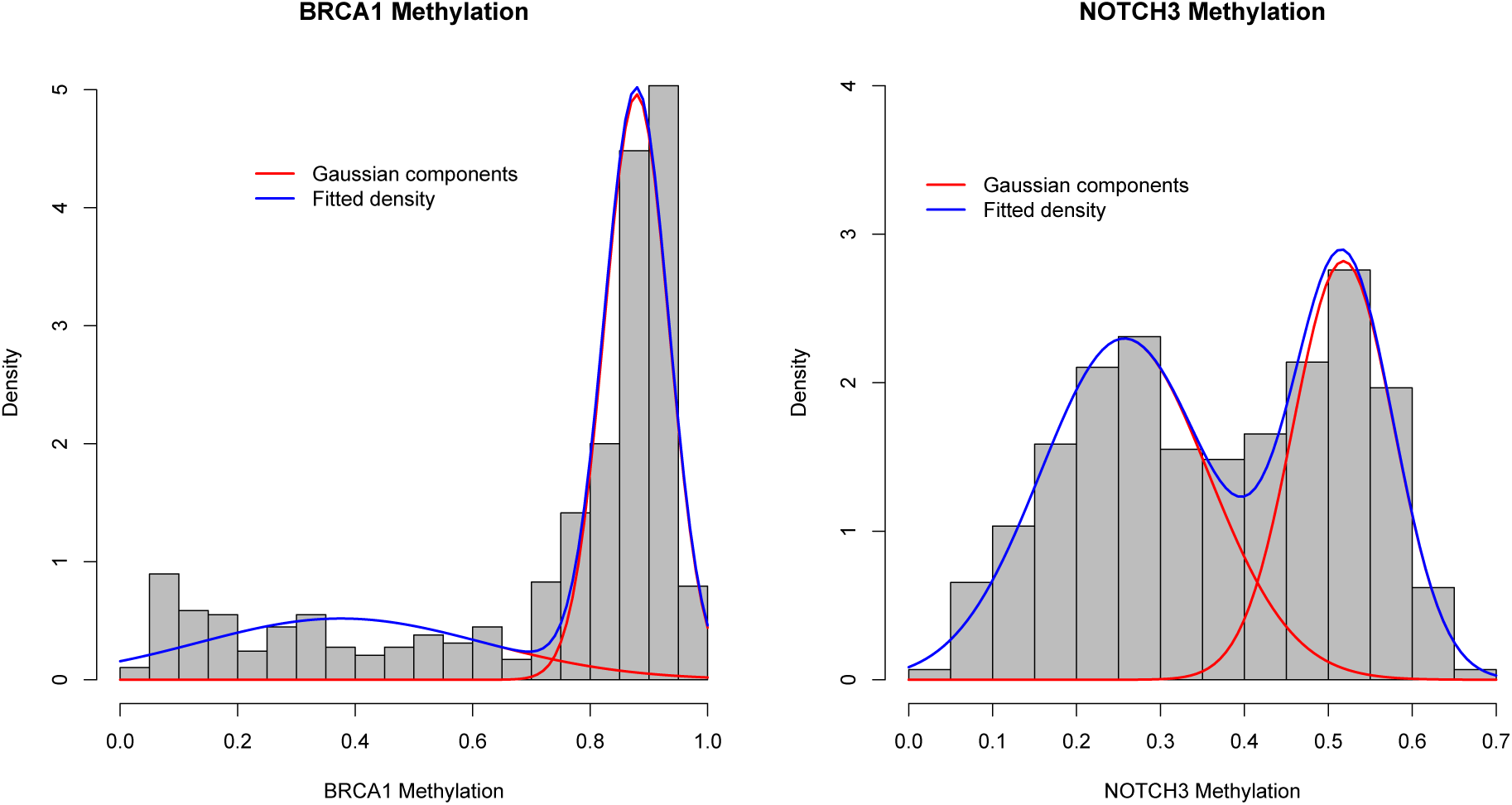
Fitted marginals by a two-component Gaussian mixture for the promoter méthylation level of gene *BRCA1* (left) and *NOTCH3* (right).

In the biological network, we assumed that the genetic or epigenetic change (CNV and DNA methylation) cannot be induced by gene expression, and imposed this constraint into our modeling (Note: this assumption is completely from biological point of view and it can be dropped without affecting our modeling and computing). The predicted graph (in Figure 7, the best-scoring network from 100 predictions) contains 73 nodes connected by 124 directed edges. Many of the edges in the graph can be confirmed in the literature. To name a few, the edge between *AURKA* and *BRCA2* may be due to the fact that a negative regulatory loop exists between *AURKA* and *BRCA2* expression in the ovarian cancer[17]. The connection between *STAT3* and *ETV6* was suggested previously that *ETV6* is a negative regulator of *STAT3* activity [18]. The edges between *RAB25* (methylation) and *RAB25* (expression) and between *CSNK2A1* (CNV) and *CSNK2A1* (expression) had been reported in several studies [10, 19, 20]. Other highly ranked edges (based on likelihood increase) include but not limited to: *STAT3*→*DLEC1*, *PTEN*→*EGFR*, *RIMBP2*→*BRCA2* and *ARID1A*→*ERD* which can be confirmed in the literature of cancer biology [10, 21, 22, 23, 24, 25]. These findings demonstrate the effectiveness of the MCBN model. In addition, as illustrated in Figure 8, the two-component Gaussian MCBN is accurate in depicting the dependency between the gene expression level and methylation level.

**Figure 7:**
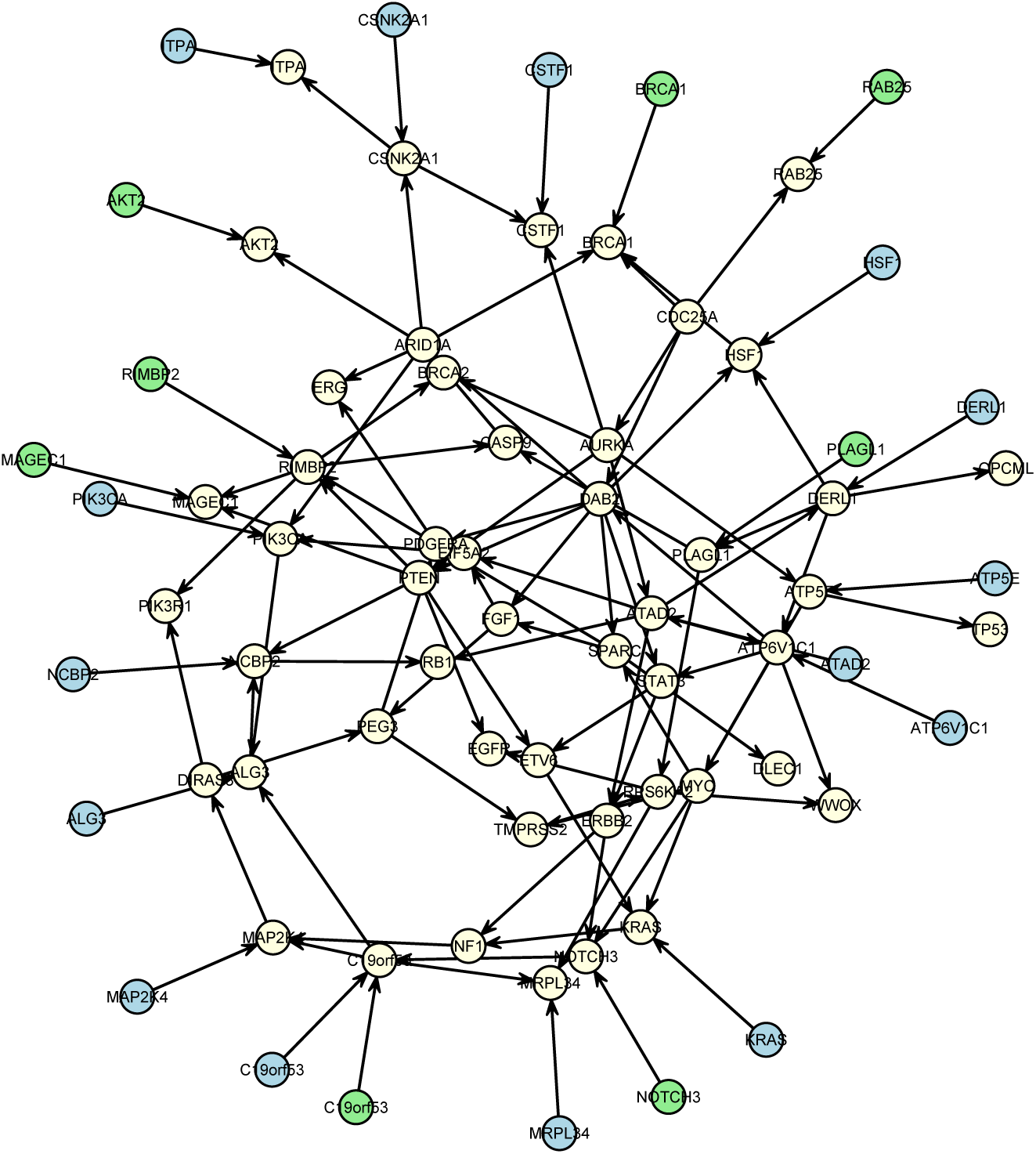
Predicted network by a two-component Gaussian MCBN model, containing the expression level of 50 genes (in light yellow), méthylation level at 8 sites (in light green) and CNV at 15 sites (in light blue).

**Figure 8:**
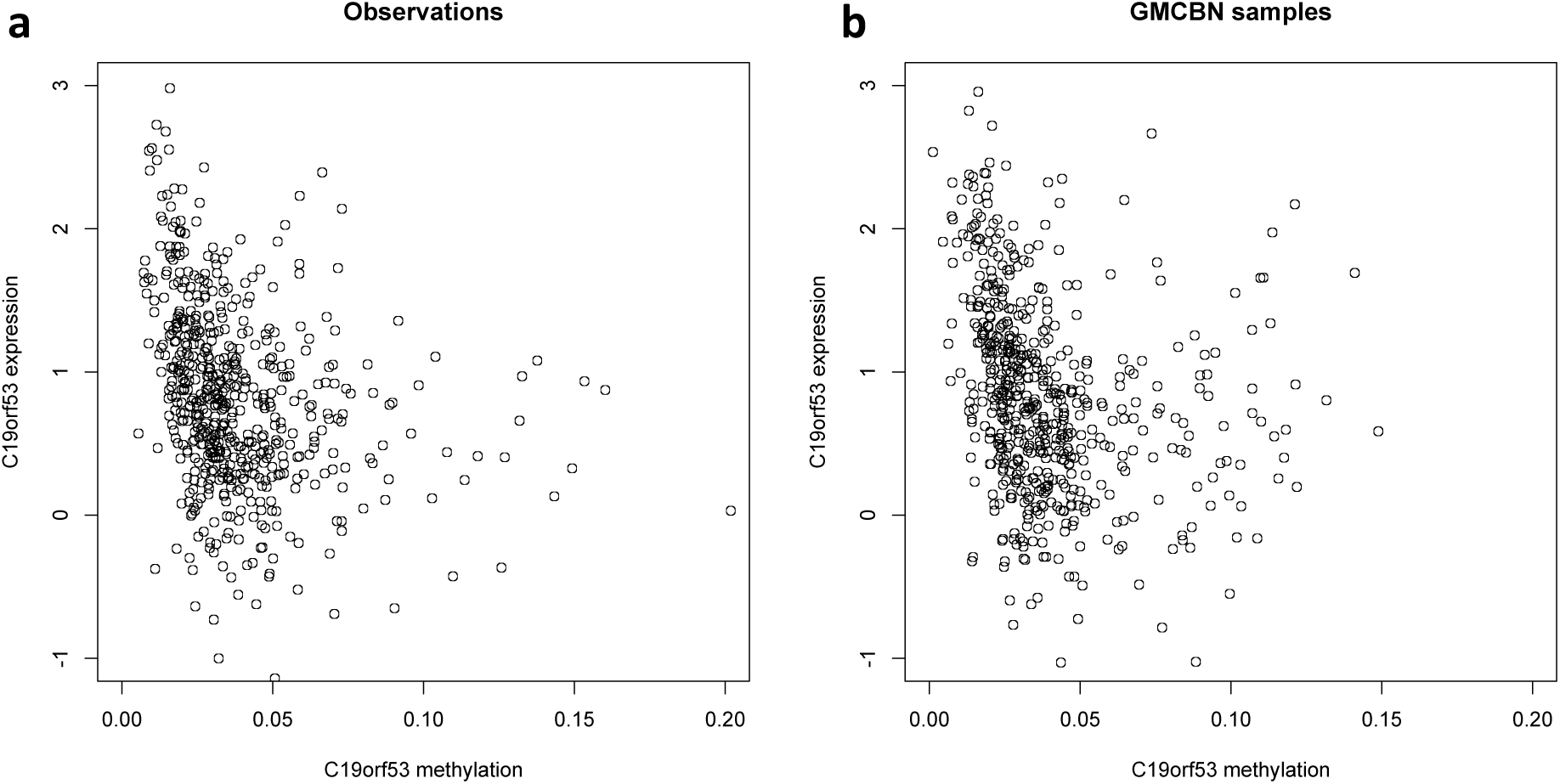
Dependence between the methylation level and expression level of gene *C19orf53:* (a) Observations; (b) Simulated samples from the two-component Gaussian MCBN.

## 6 Discussion

In this paper, we proposed a novel Bayesian network model to analyze recent cancer genomic data at the system level. The major innovation of our model is explicitly modeling the multimodal dependency structure between variables through copula function and more accurately estimating the causal network structure. The parameters in mixture copula were efficiently estimated by a routine EM algorithm, and the directed network structure was estimated by minimizing the BIC score.

The proposed Bayesian network model allows strict probabilistic inference of biological pathways, however, it also has several limitations. First, it lacks flexibility to model the cyclic mechanism due to the acyclicity constraint, for instance, A→B→A, which though may exist in gene regulatory network. Second, the parameter estimation assumes sparsity of network for computational feasibility. If the true network is dense or locally dense, the weak causations may fail to be detected. Third, due to the model complexity, the implementation of MCBN is more computationally expensive than simpler BN models such as Gaussian BN model and regular copula BN model. For large data sets, one need reduce the number of variables by filtering out irrelevant and redundant variables, and then feed the selected variables into network model for causal inference.

It is noteworthy that the Gaussian MCBN used in the two illustrative examples can be generally adapted to other mixture models such as Gamma mixture and Beta mixture. The number of mixture components can be further increased depending on the complexity of the underlying dependency structure. For relatively small data set, it is also possible to conduct statistical testing to select the best number of mixture components for each local term, however, this will significantly increase the computational complexity.

## 7 Conclusions

Understanding the biological mechanism of cancers has significant practical importance for clinical diagnosis and treatment. In this paper, we developed a mixture copula Bayesian network model for causal inference using complex cancer genomic data. The proposed model is based on finite mixture models and copula functions, and it explicitly models multimodality in the data. The graph structure and model parameters can be efficiently estimated by a routine EM approach, embedded in a heuristic search algorithm based on Bayesian information criterion. The prediction could be further improved by selecting the best-scoring model from multiple predictions with random initial values. In addition, we proposed a likelihood-based approach to select the most confident edges. The proposed MCBN model was applied to a flow cytometry data and the TCGA ovarian cancer data for inferring the causal relationships between different biological features. Compare to existing Bayesian network models, MCBN better depicts the complex dependency structure between variables, therefore may better predict the underlying causal network.

## Acknowledgement

Support has been provided in part by the Arkansas Biosciences Institute, the major research component of the Arkansas Tobacco Settlement Proceeds Act of 2000.

## Authors’ contributions

QZ conceived the study. QZ and XS analyzed the data. QZ wrote the manuscript. Both authors read and approved the final manuscript.

## Data Availability

The flow cytometry data by Sachs et al can be downloaded from http://science.sciencemag.org/content/suppl/2005/04/21/308.5721.523.DC1. TCGA ovarian cancer data can be downloaded via TCGA data portal https://tcga-data.nci.nih.gov.

## Abbreviations

TCGA: The Cancer Genome Atlas
BN: Bayesian network
GBN: Gaussian Bayesian network
CBN: Copula Bayesian network
MCBN: Mixture copula Bayesian network
EM: Expectation-Maximization
BIC: Bayesian information criterion

